# Foliar inoculation of *Phyllachora maydis* into corn induces infection and local spread in field environments

**DOI:** 10.1101/2023.12.11.571159

**Authors:** José E. Solórzano, Aarav Subbaiah, Crystal Floyd, Nathan M. Kleczewski, Dean K. Malvick

## Abstract

Tar spot of corn (*Zea mays* L.) is a significant disease in the United States and Canada caused by *Phyllachora maydis*, an obligate biotroph fungus. However, field research critical for understanding, managing, and mitigating the disease has been hindered by a need for methods to inoculate corn with *P. maydis* in field environments. In this study, we developed a method to initiate tar spot in field settings using inoculations of corn leaves with *P. maydis* inoculum that had been stored at -20 °C for 10 months. Stromata of *P. maydis* were observed 19 days after inoculation (dai), and the spread of tar spot was initially detected locally from the infection area 39 to 41 dai. Tar spot was not present in the fields beyond the inoculated area or localized spread area, signifying that the establishment of the disease resulted solely from inoculations. This study enhances our understanding of inoculation and infection of corn with *P. maydis* and tar spot development in field environments. The results will aid new research into understanding the corn tar spot pathosystem and improving management strategies.

---

*Phyllachora maydis* causes tar spot of corn (*Zea mays* L.) throughout the Americas (Valle-Torres et al. 2020) and has led to regional epidemics in the United States since 2015 that resulted in significant yield losses (Mueller et al. 2020). Colonization and development of this fungus on corn are favored by a range of environmental conditions (Hock et al. 1995; Solórzano et al. 2023a; Webster et al. 2023), but little is known regarding conditions favoring its off-season survival or conditions that favor initial infection in the field. However, the overwintered inoculum likely initiates tar spot during the growing season (Kleczewski et al. 2019; Lipps et al. 2022). Using inoculum collected from the field that had been stored at -20 °C for 5 months, infection of corn by *P. maydis* was previously achieved in controlled environments (greenhouse and growth chambers), shedding light on the survival of *P. maydis* and requirements for infection of corn (Solórzano et al. 2023a). However, despite this, a replicable methodology was needed to induce infection at larger scales in the field and to develop real-world research studies of tar spot. Therefore, the current study aimed to develop a method to induce *P. maydis* infection into corn and observe its establishment and development in field settings.

The inoculum used for this study was arbitrarily selected, tar spot-infected corn leaves with > 20% disease severity (Fig. 1A; I) collected in September 2021 and 2022 from corn fields in southern Minnesota in the region where tar spot has occurred since 2019 (Malvick et al. 2020). The Infected-senesced leaves collected in 2021 (Source I) were placed in cloth sacks and stored at 25 ± 1 °C until use. Infected leaves collected in September 2022 from actively growing plants (Source II) were placed in plastic bags and stored at -20 °C until use (Solórzano et al. 2023a). The leaves from either source were not surface sterilized before or following storage.

**Figure 1.**
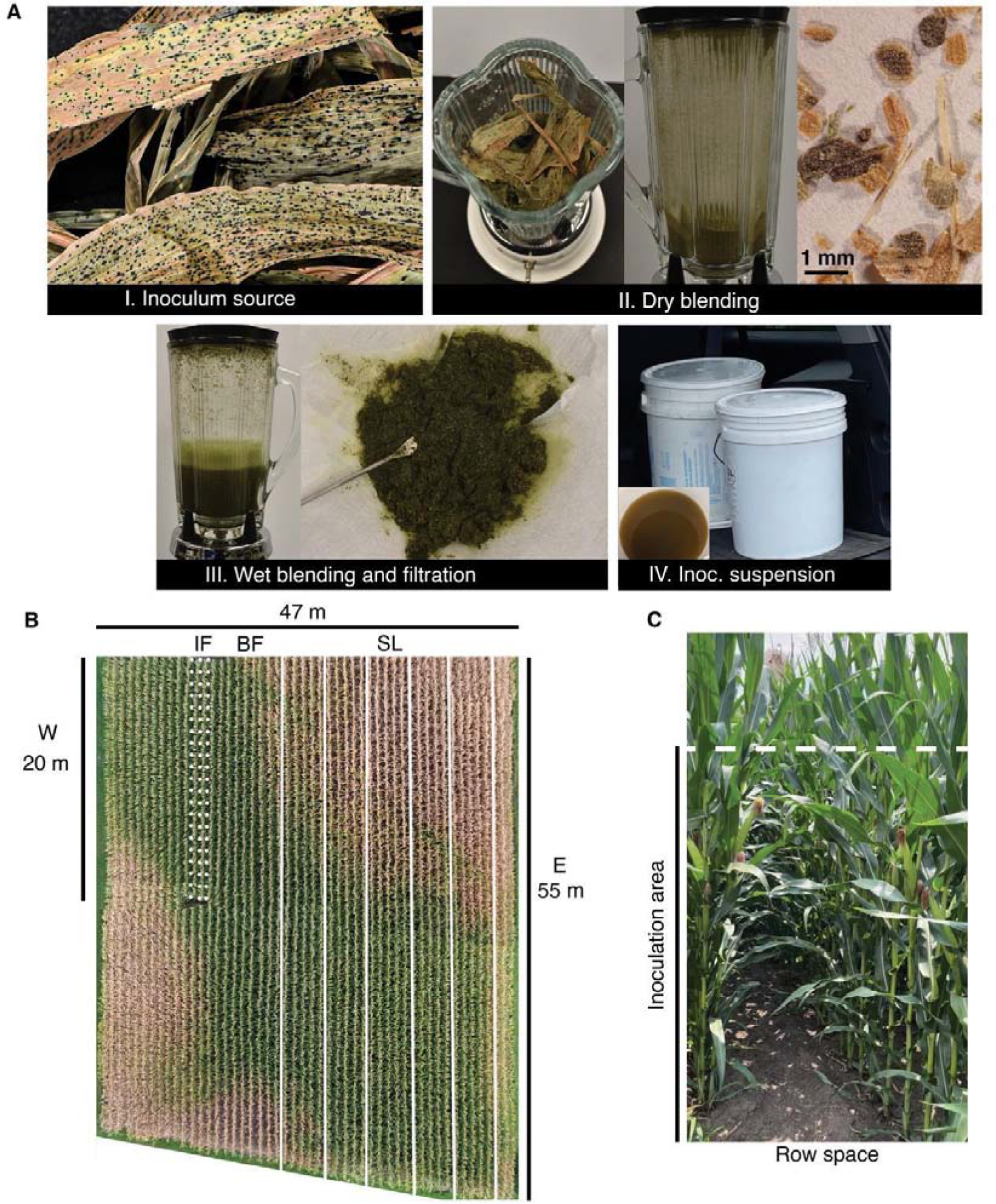
Illustrations of the procedure to generate inoculum suspensions and depiction of the main field area used for inoculation. **A**. Method used to extract spores from stromata of *Phyllachora maydis* (I), dry blending of infected corn leaves to break stromata into fine particles (II), wet blending and filtration to extract spores (III), and storage and transportation of inoculum suspension (IV). **B**. Corn research plot at the University of Minnesota, Saint Paul, photographed 45 days following the inoculations, where the brown color indicates advanced stages of drought stress in parts of the field. W is the western section of the research plot. The infection focus (IF) is where Source I derived inoculum was applied, denoted by dotted lines. The buffer area (BF) is the non-inoculated area (5.3 m) between inoculated rows (IF and SL). Inoculum suspension from Source I was applied to corn plants (denoted SL) to one row to the left and one to the right of each white line. **C**. Plants’ area/foliage targeted for inoculation delimited by the dashed lines.

The presence of ascospores in stromata and their germination were assessed before preparing inoculum suspensions (Solórzano et al. 2023a, b), and leaves from Source II were allowed to thaw at 25 ± 1 °C for 24 h to aid in breaking open the stromata. Stromata were fragmented into small pieces, and spores were extracted following the methods described by Solórzano et al. (2023a), adapted for this study. The midrib of each leaf was manually removed, and the remaining leaf tissue was ground into fine particles with an industrial blender (Waring 700G Single-Speed Blender, Model WF2212112) (Fig. 1A; II). Then, to extract spores, the ground leaf tissue and stomata were mixed with autoclaved deionized water at a 1:3 ratio (w/v) and blended for 1 min (Fig. 1A; III). This process was conducted in small batches to increase the chances of extracting spores. Spores were collected by filtrating this mixture (Fig. 1A; III) through 4 layers of cheesecloth and using a metal spatula to press and revolve the mixture (Kleczewski et al. 2019). The spore concentration was determined with a hemacytometer and adjusted to 10^4^ ascospores mL^-1^ in 0.01% Tween 20 (Solórzano et al. 2023a). One batch of inoculum suspension (56 L) was prepared from the 22-month-stored Source I leaves (n = 60). Another batch of inoculum suspension (13 L) was prepared from the 10-month-stored Source II leaves (n = 9). Inoculum suspensions were maintained at 21 °C during transport to the research plot and used < 2 h after filtration (Fig. 1A; IV).

For inoculations, the corn hybrid GC-103-58 RSS (Gold Country Seed^®^), known to be susceptible to *P. maydis* (Solórzano et al. 2023a), was planted on May 4, 2023, with 16 cm seed spacing and 81 cm row spacing in a field with a previous soybean-barley-corn rotation at the University of Minnesota Saint Paul research plots. This research area was selected because, in 2021 and 2022, tar spot occurred naturally in neighboring plots at low plant incidence and severity < 1%. The plot (Fig. 1B) was fertilized with urea at 200 kg nitrogen ha^-1^ before planting and 45 kg ha^-1^ of ammonium sulfate 35 days after planting. The herbicides Harness® at 2 L ha^-1^ and Hornet WDG at 0.3 L ha^-1^ were applied at planting, and glyphosate Cornerstone® 5 Plus at 2 L ha^-1^ was applied 27 days after planting. Plants were not sprayed with fungicides. The site relied on natural rainfall, and supplied irrigation was not used. The same corn hybrid was planted on May 5 and managed following the same cropping conditions as described above in a secondary research site located 800 m from the main site described above.

Plants at reproductive growth stage 2 (R2) were inoculated at 2:00 p.m. on July 22, 2023 (Fig. 2A). The inoculation date was selected based on the plants’ growth stage (R2) and the observed 2022 tar spot initial occurrence date in southern Minnesota minus 15 days, representing the maximum incubation period recorded during epidemiological studies of tar spot in the field (Hock et al. 1995). At the time of the inoculation, the corn leaves and soil surface were dry in and near the inoculated plots, and tar spot stromata were absent in the plot, as well as in other corn in the area. During inoculation, the relative humidity (RH) was 35%, the dew point temperature was 15 °C, and the air temperature was 28 °C (Fig. 3). Inoculum suspensions were sprayed to completely cover the adaxial surface of the leaves using an electric-powered sprayer (RYOBI ONE+ 18V 1 Gal. Chemical Sprayer, Model P2800; 440 mL min^-1^), primarily targeting the leaves (9 to 10) below and one leaf collar above the ear leaf (Fig. 1C). Care was taken to avoid disturbing the inoculated leaves by walking backward through the plot during application of the inoculum. Each inoculum suspension type was applied to different areas of the research plots. The inoculum suspension from Source I was applied to plants (n ∼ 3000) in rows on the east side of the main corn plot (Fig. 1B; SL), leaving 5.3 m (Fig. 1B; BF) of non-inoculated plants as a control group. The inoculum suspension prepared from Source II was applied to a set of plants (n = 592) on the west side of the plot area, designated as the infection focus (IF) (Fig. 1B; IF). The remaining inoculum from Source II (∼ 1 L) was applied to the ear leaf of 130 plants on the southern border of the secondary research plot, covering between 15% to 20% of the leaf surfaces. Each inoculated plant was an experimental unit. Shortly after inoculations (∼ 5 min), light rain (0.7 mm) fell for < 30 min, resulting in left wetness, 75% to 80% RH, dew point temperature of 14 to 15 °C, and air temperature of 19 °C. The rain did not continue that day. The night after inoculation, RH was 78% to 86%, and the air temperature was 15 °C to 18 °C. The dew point temperatures reached a maximum of 15 °C and a minimum of 12 °C (Fig. 3). The next rain event (7.6 mm) occurred 2 days after the inoculations. The largest rainfall event (45 mm) was reported at night, 12 days after inoculations (Fig. 3).

**Figure 2.**
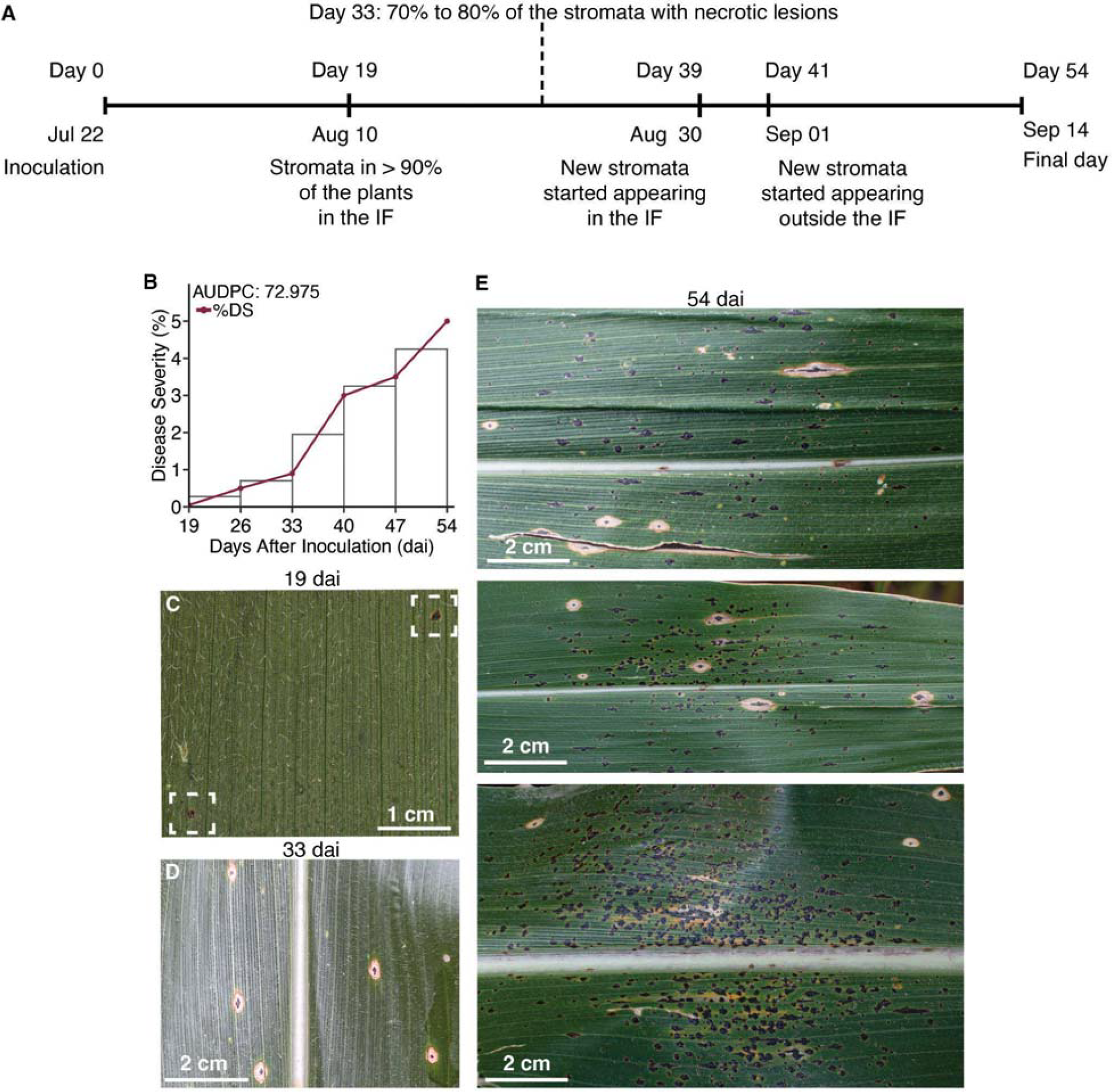
Foliar inoculation of corn (*Zea mays*) leaves with *Phyllachora maydis* led to tar spot establishment and development in field conditions. **A**. Timeline for induction of *P. maydis* infection into corn, stroma development, and dispersal in and outside the infection focus (Fig. 1B; IF). **B**. Progress of tar spot severity based on leaf area coverage and represented as the area under the disease progress curve (AUDPC). **C**. Emerging stromata, 19 days after inoculation (dai), denoted by dashed squares. **D**. At 33 dai, 70 to 80% of stromata that had emerged at 19 dai were surrounded with necrotic lesions. **E**. Representation of tar spot severity with old and new stromata 54 dai. Stromata that started to appear in the second cycle (39 dai) are those without necrotic lesions.

**Figure 3.**
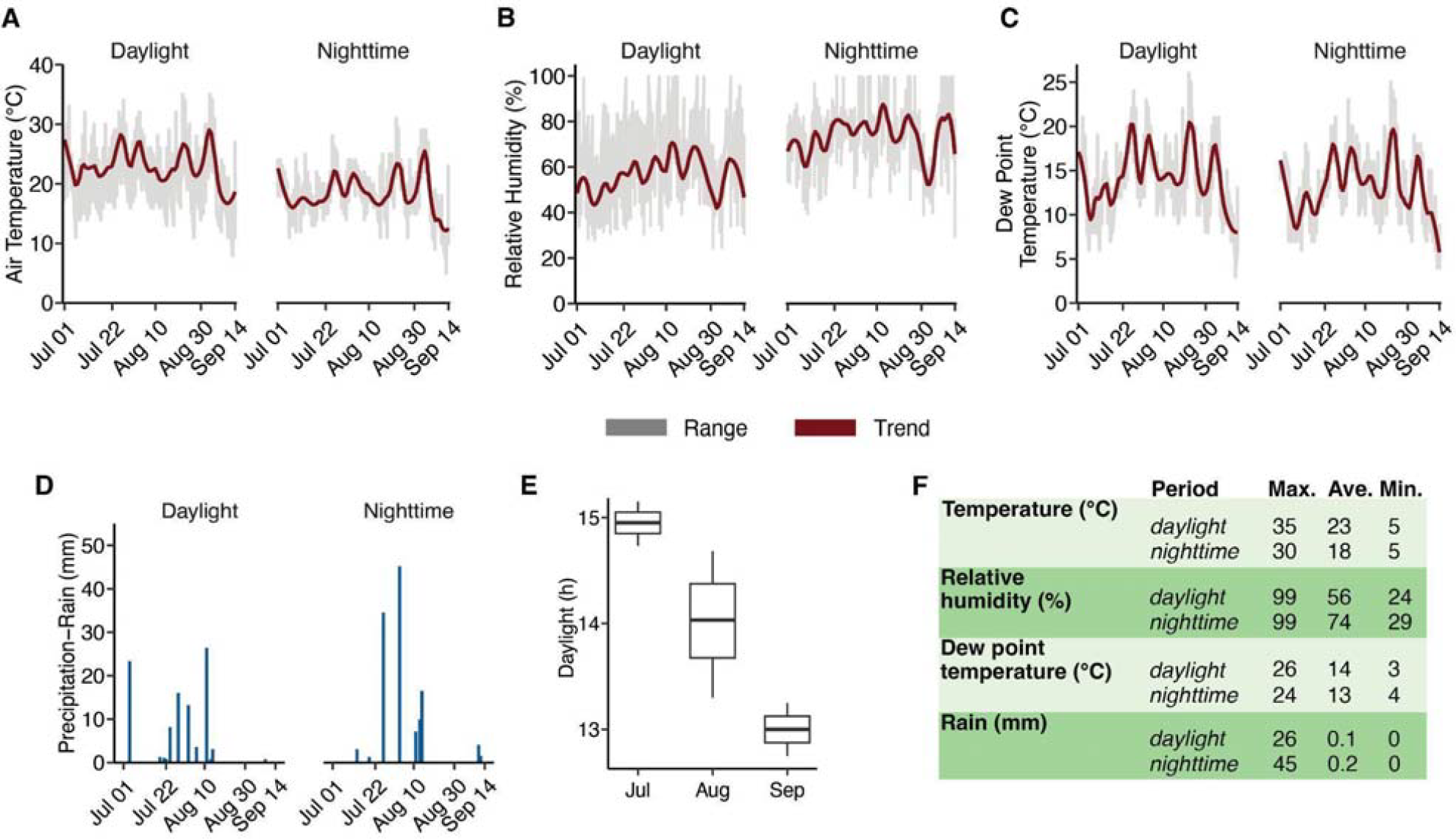
Weather conditions before, during, and after inoculating corn leaves with *P. maydis* on July 22 led to the initiation and development of tar spot. **A to D**. Day and night variability of weather conditions at the research area. Data shown for A to C are the hourly daily range (gray) and the trend variability of the data calculated using a locally weighted least squares regression (maroon). **D**. Precipitation (rain) records (mm) for days and nights. **E**. Daylight duration (h) per month from July 1 to September 14, the final day of the study. **F**. Summary of weather data gathered during this study from July 1 to September 14, where Max. is maximum, Ave. is average, and Min. is minimum.

*P. maydis* infected corn leaves in both research plots after applying the inoculum suspension prepared from Source II. Stromata were observed 19 days after inoculation (dai) and were present in > 90% of the inoculated plants in the IF area of the main research site and in < 10 % of the plants inoculated in the secondary research site. (Fig. 2A to 2C). Tar spot was identified as described by Solórzano et al. (2023b). This period of 19 days to stromata formation is longer than the 10 to 12 days observed for this corn hybrid following inoculations at the vegetative growth stage 2 (V2) in controlled environments (Solórzano et al. 2023a). However, the reasons for the differences in time periods for the formation of stromata are unknown but may have resulted from the interplay of the pathogen and host, moderated by the plant’s growth stage and the environment. At 19 dai in the field, stromata were absent in non-inoculated plants, plants inoculated with inoculum suspension from Source I, and plants in the adjacent corn fields. Within 6 days (25 dai), all the inoculated plants in the IF of the main site were infected, each containing multiple stromata of *P. maydis* in different leaves. In addition, at this time, > 50% of stromata also became surrounded by necrotic lesions (fisheye) that were visible without magnification, similar to what Hock et al. (1992) reported. However, necrotic lesions were not observed around the stromata in the plants inoculated in the secondary research site. By 33 dai, the necrotic lesions had developed around 70% to 80% of the stromata on the plants and were present on > 95% of the inoculated plants in the IF in the main research site (Fig. 2D and 2E). The presence and number of necrotic lesions observed were unexpected since the inoculum source contained mostly stromata without necrotic lesions (Fig. 1A; I). The etiology of necrotic lesions around the stromata is fundamentally unknown; however, recent studies indicate that various fungi may be associated with the lesions (Luis et al. 2023; McCoy et al. 2019).

This study presents for the first time a precise assessment of the latent period of *P. maydis* (time from inoculation to reproduction and secondary infectivity) in the field, determined to be between 20 to 22 days and its secondary spread. The second cycle of tar spot was observed 39 to 41 dai, when new and small stromata emerged on the inoculated plants in the IF and plants beyond this infected area, respectively (Fig. 2A). In contrast, this was not observed in the infected plants in the secondary research site where only 1 or 2 stromata formed in each plant. The low incidence and severity of tar spot on those plants is likely due to a lower quantity of inoculum applied compared to inoculations conducted in the IF of the main research site.

The newly emerging stromata on the plants in the IF area of the main research site were found aggregated on the leaves, usually surrounding the older stromata (Fig. 2A and 2E). Outside the IF, infected plants increased in frequency 42 to 46 dai and were initially in neighboring rows to the west, east, and south. The number of plants with stromata decreased with distance from the IF, and the incidence and severity were low, i.e., one stroma on a single leaf in some plants (Fig. 1B). At that time, necrotic lesions were not observed around the new stromata as observed in the first cycle (Fig. 2E). The infected plants outside the IF were at a maximum distance of 5.7 m west, 14.2 m east, and 4 m south. In addition to the previous emergence of stromata, more continued appearing in plants in the IF after the second cycle was initiated (Fig. 2E), possibly due to a varied timing of the release of spores from the stromata that formed during the first cycle, 19 to 25 dai. At 54 dai, disease severity in the IF reached an average of 5% based on leaf area coverage (Fig. 2B), and severity was < 1% outside the IF. In addition, stromata were seen for the first time at 54 dai in border plants (n = 9) in an adjacent corn research plot 4.5 m from the inoculated plants. This local spread likely was favored by wind because anthropochory was avoided by not entering areas other than the IF for 17 days after stromata were first seen 19 dai. Altogether, by applying the methodology developed in this study, we knew exactly when and where plants were exposed to *P. maydis* inoculum and when stroma formed and developed in the absence of any natural infection.

This study highlights the importance of incorporating surveillance of *P. maydis* in epidemiological studies of tar spot. It also emphasizes the importance of inoculum availability, viability, and/or amount, as it was clear in this study that the host and environment (Hock et al. 1995; Solórzano et al. 2023a; Webster et al. 2023) were compatible with tar spot occurrence and that only the inoculum was needed to initiate the disease. During this study, tar spot did not occur naturally in any corn beyond the research area during the growing season, arguably due to the absence of inoculum or viable inoculum and indicating that environmental conditions alone do not predict tar spot occurrence. Weather conditions varied before and after inoculation (Fig. 3), favoring the establishment and development of tar spot. For example, before and during the study, the air temperature varied from 18 to 28 °C (Fig. 3), similar to the reported ranges by Hock et al. (1995) and by Solórzano et al. (2023a) and as recently discussed by Webster et al. (2023). The period following inoculations was generally dry with periodic rain over the first 19 days and then dry, but stroma and tar spot continued developing. The day and night mean RH was 56% to 74%, respectively (Fig. 3). However, the RH exceeded 90% on different non-consecutive days, 7 days preceding stromata formation, and 15 days after that, which is a low frequency of occurrence (< 25%) compared to the frequency of the observed mean %RH (Fig. 3). Thus, frequent rainfall and high moisture were not required for tar spot to develop, which contradicts the information on high moisture conditions favoring tar spot development as reported by Hock et al. (1995). Solórzano et al. (2023a) previously reported that *P. maydis* can infect and develop at low %RH in controlled environments after a short (20 h) period in 100% RH in darkness. This is similar to the events that occurred after inoculation in the current field study, where inoculation followed by light rain resulted in the establishment of the disease (Fig 3).

In conclusion, a single inoculation of *P. maydis* into corn leaves at an early corn reproductive growth stage led to colonization, infection, and spread of *P. maydis* and the development of tar spot under variable conditions in field environments (Fig. 3). This is the first report to our knowledge of successful foliar inoculation of *P. maydis* in field settings that resulted in infection, development, and spread of tar spot. The current study presents a method for inducing *P. maydis* infection into corn in the field and details information on the storage and preparation of *P. maydis* inoculum and methods to conduct foliar inoculations. This short communication also provides information on environmental conditions favorable for tar spot occurrence and supports additional studies on *P. maydis* and the disease.

## Acknowledgment

We thank the Minnesota Invasive Terrestrial Plants and Pests Center for supporting this research, and we thank Peter Boulay, Julian Cooper, and field technicians at the University of Minnesota for assistance with the study.

## Literature cited

Hock, J., Dittrich, U., Renfro, B. L., and Kranz, J. 1992. Sequential development of pathogens in the maize tarspot disease complex. Mycopathologia. 117:157–161.

Hock, J., Kranz, J., and Renfro, B. L. 1995. Studies on the epidemiology of the tar spot disease complex of maize in Mexico. Plant Pathol. 44:490–502.

Kleczewski, N. M., Donnelly, J., and Higgins, R. 2019. Phyllachora maydis, causal agent of tar spot on corn, can overwinter in northern Illinois. Plant Health Prog. 20:178–178.

Lipps, S., Smith, D., Telenko, D., Paul, P., Kleczewski, N., and Jamann, T. 2022. Identification of resistance for Phyllachora maydis of maize in exotic-derived germplasm. Crop Sci. 62:859–866.

Luis, J. M., Mehl, H. L., Plewa, D., and Kleczewski, N. M. 2023. Is Microdochium maydis associated with necrotic lesions in the tar spot disease complex? A culture-based survey of maize in Mexico and the Midwestern United States. Phytopathology®. 113:1890–1897.

Malvick, D. K., Plewa, D. E., Lara, D., Kleczewski, N. M., Floyd, C. M., and Arenz, B. E. 2020. First report of tar spot of corn caused by Phyllachora maydis in Minnesota. Plant Dis. 104:1865–1865.

McCoy, A. G., Roth, M. G., Shay, R., Noel, Z. A., Jayawardana, M. A., Longley, R. W., Bonito, G., and Chilvers, M. I. 2019. Identification of fungal communities within the tar spot complex of corn in Michigan via next-generation sequencing. Phytobiomes J. 3:235–243.

Mueller, D. S., Wise, K. A., Sisson, A. J., Allen, T. W., Bergstrom, G. C., Bissonnette, K. M., Bradley, C. A., Byamukama, E., Chilvers, M. I., Collins, A. A., Esker, P. D., Faske, T. R., Friskop, A. J., Hagan, A. K., Heiniger, R. W., Hollier, C. A., Isakeit, T., Jackson-Ziems, T. A., Jardine, D. J., Kelly, H. M., Kleczewski, N. M., Koehler, A. M., Koenning, S. R., Malvick, D. K., Mehl, H. L., Meyer, R. F., Paul, P. A., Peltier, A. J., Price, P. P., Robertson, A. E., Roth, G. W., Sikora, E. J., Smith, D. L., Tande, C. A., Telenko, D. E. P., Tenuta, A. U., Thiessen, L. D., Wiebold. W. J. 2020. Corn yield loss estimates due to diseases in the United States and Ontario, Canada, from 2016 to 2019. Plant Health Prog. 21:238–247.

Solórzano, J. E., Issendorf, S. E., Drott, M. T., Check, J. C., Roggenkamp, E. M., Cruz, C. D., Kleczewski, N. M., Gongóra-Canul C. C., and Malvick, D. K. 2023a. A new and effective method to induce infection of Phyllachora maydis into corn for tar spot studies in controlled environments. Plant Methods. 19:83.

Solórzano, J. E., Cruz, C. D., Arenz, B. E., Malvick, D. K., and Kleczewski, N. M. 2023b. Tar spot of corn: A diagnostic and methods guide. Plant Health Prog. 24:117–122.

Valle-Torres, J., Ross, T. J., Plewa, D., Avellaneda, M. C., Check, J., Chilvers, M. I., Cruz, A. P., Dalla Lana, F., Groves, C., Gongóra-Canul, C., Henriquez-Dole, T., Jamann, T., Kleczewski, N., Lipps, S., Malvick, D., McCoy, A. G., Mueller, D. S., Paul, P. A., Puerto, C., Schloemer, C., Raid, R. N., Robertson, A., Roggenkamp, E. M., Smith, D. L., Telenko, D. E. P., and Cruz, C. D. 2020. Tar spot: An understudied disease threatening corn production in the Americas. Plant Dis. 104:2541–2550.

Webster, R. W., Nicolli, C., Allen, T. W., Bish, M. D., Bissonnette, K., Check, J. C., Chilvers, M. I., Duffeck, M. R., Kleczewski, N., Luis, J. M., Muller, B. D., Paul, P. A., Price, P. P., Robertson, A. E., Ross, T. J., Schmidt, C., Schmidt, R., Schmidt, T., Shim, S., Telenko, D. E. P., Wise, K., Smith, D. L. 2023. Uncovering the environmental conditions required for Phyllachora maydis infection and tar spot development on corn in the United States for use as predictive models for future epidemics. Sci. Rep. 13:17064.

